# Climate change and the migration of a pastoralist people c.3500 cal. years BP inferred from palaeofire and lipid biomarker records in the montane Western Ghats, India

**DOI:** 10.1101/2020.08.15.252189

**Authors:** Sarath Pullyottum Kavil, Prabhakaran Ramya Bala, Pankaj Kumar, Devanita Ghosh, Raman Sukumar

## Abstract

Human migration in response to climate change during the Holocene has been recorded in many regions of the world. The Todas are a pastoralist people who are believed to have colonized the higher elevations (>2000 m asl) of the Nilgiris in the Western Ghats, India, not earlier than about 2000 cal. yr BP. Vegetation shifts in response to changing climate in tropical montane forest-grassland mosaic of the Ghats have been well documented using stable carbon isotopes and pollen profiles; however, there have been no corresponding investigations of human presence and activity at the highest elevations. We used a number of other proxies to infer the human ecology of this montane region. Radiocarbon dated (~22,000 cal. BP to the present) peat samples from the Sandynallah basin (2200m asl, Nilgiri hills, Tamil Nadu State) were used to reconstruct fire history, animal abundance, and human presence since the Last Glacial Maximum (LGM). While the macro-charcoal record indicates fires at the LGM, macro- and micro-charcoal counts indicate intense fire at ~3500 cal. yr BP, coprophilous fungal spores indicate a large population of herbivorous mammals, and steroid biomarkers indicate human faecal remains for the first time. This period is also characterized by dry arid conditions and dominant grassy vegetation as inferred from *n*-alkane signatures. We thus infer that a pastoralist people, most likely the Todas, migrated to the highest elevations of the Western Ghats along with their buffalo herds in response to prolonged or abrupt climate change in peninsular India, about 3500 cal. yr BP or at least 1500 years prior to what historical accounts assume.

## 1. Introduction

Climate change is known to have driven human adaption through migration from the earliest times of human evolution (Carto et al. 2009; Tierney, deMenocal, and Zander 2017) through to the historical period of the late Holocene (D’Andrea et al. 2011). Some of the best documented migrations involving nomadic pastoralists and agriculturalists come from Chinese chronicles (Fang and Liu 1992; Pei 2017). A series of migrations of nomadic people in the Mongolian grasslands and east-central Asia between 190 BCE and 1880 CE is closely related to historical cold and dry periods (Fang and Liu 1992). Holocene migrations in the tropics have been less well documented, though the migration of the Harappan people southward, following the decline of the extensive Harappan civilization in northwest India and Pakistan, has been increasingly attributed to the aridification of this region about four millennia ago (Chandran, 1997; Dixit, Hodell, and Petrie 2014; Dutt et al. 2019).

Peninsular India has been inhabited by modern humans not long after the out-of-Africa migration (Korisettar 2016) while, at the same time, it has been subject to a changing climate in tune with global change during the late Pleistocene and Holocene Epochs (Sukumar et al. 1993; Rajagopalan et al., 1997; Bhagwat, Nogué, and Willis 2012). Human societies in the peninsula have gone through a complex process of transitions from hunter-gatherer through pastoralist to settled agricultural over this period (Korisettar et al. 2001; Johansen 2004; Fuller, Boivin, and Korisettar 2007; Fuller 2013; Morrison et al. in press), though their links to climate have been scarcely investigated. The Western Ghats (sea level up to >2600m in the Nilgiris and Anamalais) running parallel to the west coast of India have generally been settled much after the Deccan plateau and its river valleys, perhaps because the Ghats were largely hilly and densely forested. Although human occupation of the Ghats has only been attributed to the end of the Palaeolithic, about 12,000 cal. BP (Chandran 1997), it is certain that human influences along the fringes of the Ghats, both the moister west and the drier east, would have been much earlier. In any case, no evidence has so far been presented or even suggested for human occupation at the highest elevations (>2000m asl) of the Ghats prior to about 2000 cal. BP; the steep slopes, rugged terrain and dense forests at lower to mid-elevation with a history of malaria may have proved to be difficult barriers for the ascent of humans to the highest elevations. The Todas, an obligate buffalo-tending pastoral community, are widely believed to be the earliest people to move into the upper Nilgiri plateau perhaps during the 1st millennium CE or later (Rivers 1906; Noble 1976; Emeneau 1997; Zagarell 1997). Other indigenous people of the Nilgiris include the artisan Kotas, and the hunter-gather or cultivator Kurumbas and Irulas who have inhabited the region since early times, while the cultivator Badagas were 16th century immigrants (Hockings 1980).

The higher elevation (>2000m asl) of the Nilgiris features a unique ecosystem mosaic consisting of patches of tropical montane forests (locally known as *sholas*) restricted to the folds, valleys and depressions, and extensive grasslands on the slopes, ridges and exposed areas (Meher-Homji 1967; Jose et al. 1994; Das et al. 2015). Late Pleistocene and Holocene paleoclimatic reconstruction using stable carbon isotope analysis of peat from high-elevation (>2000 m asl) valleys of the Nilgiris suggests that the region had undergone phases of wet and arid climate, broadly in tune with well-known global climatic events since the Last Glacial Maximum, resulting in shifts in the relative dominance of C3 plants (mainly woody vegetation and herbs other than tropical grasses) and C4 plants (tropical grasses) (Sukumar et al. 1993; Caner et al. 2007). Evidence from pollen studies also suggests the co-occurrence of grassland and forest through moister and more arid phases in the Nilgiri region over this period (Vasanthy 1988; Sutra, Bonnefille, and Fontugne 1997). While these studies have provided us with a fairly good understanding of past vegetation and associated climate, in particular the strength of the monsoon, they have not specifically investigated the human ecology associated with these changes.

At the same time, several other proxies such as macro- and microcharcoal abundance in sediments to record fires including those of possible anthropogenic origin (Carcaillet et al. 2001; Whitlock and Larsen 2002; Clark 1988), fungal spores associated with dung to assess animal abundance (Van Geel 2002), leaf wax lipids characteristic of different plant types, and steroid compounds indicative of animal- and human faeces (Pancost et al. 2002; Meyers 2003) are useful to understand the relationship between climate, vegetation and human presence or activity. In particular, lipid biomarkers which are powerful tools in resolving past plant or animal origins have been hardly explored in paleoecological research in India. Alkanes have been used to trace their source in sediments from microbes, algae and higher plant types (Cranwell 1973; Ficken et al. 2000; Pancost et al. 2002; Nichols et al. 2006). *n*-alkane distributions dominated by carbon chain lengths C_21_, C_23_ and C_25_ suggest aquatic plant dominance (Pancost et al. 2002; Nichols et al. 2006). A predominance of C_27_, C_29_ and C_31_ indicate higher contribution from epicuticular leaf waxes of land plants, with elevated C_31_ presence attributed to inputs from grasses and C_27_ or C_29_ predominance related to a tree-leaf origin (Cranwell 1973; Rommerskirchen et al. 2006). Thus, *n*C_27_/*n*C_31_ ratio has been applied to understand the changes in relative distributions of grasses and woody plants in peatland (Schwark, Zink, and Lechterbeck 2002).

Faecal steroid (stanols and sterols) biomarkers have been used to identify the biogenic origin in ancient samples as these organic molecules are resistant to diagenetic alteration and degradation (Bull et al., 1999; Bull et al., 2002; Prost et al., 2017). Stanols are produced by the microbial degradation process from their Δ^5^-sterols precursors (Bull et al. 2002; Nash et al. 2005). Coprostanol is the major 5β-stanol in human faeces, while the faeces of herbivores such as cows and sheep contain a higher relative proportion of 5β-stigmastanol due to the high amount of stigmasterol, campesterol and sitosterol derived from their herbivorous diet (Linseele et al. 2013; Leeming et al. 1996). The ratios of various steroid compounds in dated archaeological samples have thus been used in identifying human and herbivorous mammalian faeces and, hence, their presence at a particular site (Linseele et al. 2013; Prost et al. 2017; Schroeter et al. 2020). Steroid biomarkers have never been used so far in exploring their origins in any archaeological or paleoecological context in India.

We thus carried out this study on Late Quaternary paleoecology of the Nilgiris with the following primary objectives:

a. To complement the earlier studies on paleo-climatic and paleo-vegetation by using proxies other than pollen and stable isotopes.
b. To use charcoal counts in peat to trace fire history and draw inferences on its possible relationship to past climate and human activity.
c. To use fungal spores specific to animal dung to determine the past abundance of herbivorous animals which may also signify presence of domestic animals.
d. To use lipid steroid biomarkers (associated with faeces) extracted from peat to detect the presence of domestic livestock and human populations in the Nilgiri plateau.

## 2. Methods

### 2.1. Study site

Our study site, the Sandynallah valley, is located between 11°26’32”N, 76°38’6”E and 11°26’37”N 76°38’8”E at an elevation of ~2200 m above sea level, in the Nilgiri massif of the southern Western Ghats, India (Fig. 1). Since 1950, the study site is administered by the Sheep Breeding Research Station (SBRS) of Tamil Nadu University of Veterinary and Animal Sciences (TANUVAS). Although the site is located in the tropical belt, peat is preserved in valleys in this mountain range due to cool temperatures (annual average 13.5 °C) and moderately high precipitation leading to water logging (annual average precipitation of about 1400 mm) (von Lengerke 1977). These tropical mountain ranges feature peat deposits dating back to >40,000 radiocarbon yr BP which have been studied extensively to reconstruct Late Quaternary vegetation and climate (Vasanthy 1988; Sukumar et al. 1993; Sutra, Bonnefille, and Fontugne 1997; Rajagopalan et al., 1997; Ramya Bala 2015; Ramya Bala et al. 2016).

**Fig 1.**
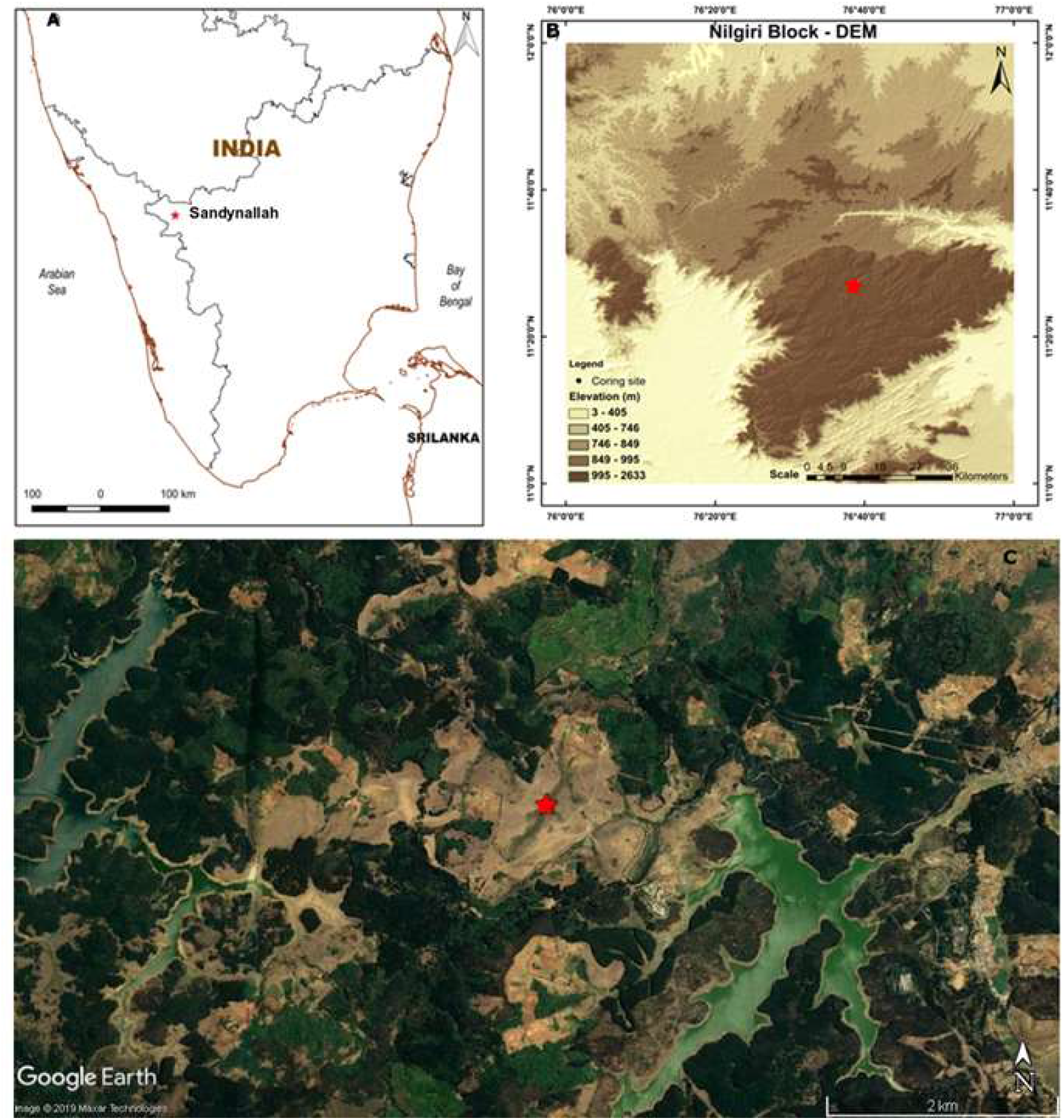
(A) The physical map of peninsular India depicting the Sandynallah site in the Nilgiris (B) The digital elevation model (source: ASTER GDEM) of the study region with the altitude range given in the side panel, (C) Google Earth image of Sandynallah and the trenching site with a mosaic vegetation pattern.

### 2.2. Sample collection and ^14^C dating

A pit of ~1.8 m depth was dug close to a location from which a core (labelled as Core 1) had been previously taken and radiocarbon dated (Ramya Bala et al. 2016), and peat samples collected as monoliths at regular intervals in a stainless-steel box of dimensions 15.6 cm x 9.3 cm x 3.0 cm. Thirteen samples were collected in zip lock bags, adjacent samples starting from the surface separated by ~2 cm, down to a depth of ~1.6 m. Sub-samples were freeze dried (Labconco bulk drier) and crushed into fine powder and stored. Accelerated Mass Spectrometry (AMS) radiocarbon dating of seven peat samples after standard Acid Alkali Acid (AAA) pre-treatment (Nakamura et al., 2003) was carried out at the Inter-University Accelerator Centre (IUAC), New Delhi (Table 1). Blank sample prepared along with the peat samples was used for background correction, while standard sample OXII (oxalic acid II) was used for normalization (Sharma et al., 2019). Data quality was monitored with a secondary standard (IAEA-C7); the consensus value (pMC = 49.53±0.12) was within the error limits of the experimental result (pMC= 49.48±0.12). Calibration of the radiocarbon dates was done using OxCal (Ramsey, 2013).

**Table 1.** Details of the AMS ^14^C dated peat samples and calibrated dates using OxCal from the Sandynallah basin

| Lab Code | Sample type | Sample ID | Depth range | AMS $^{14}\text{C}$ age ± Error | Calibrated age range Cal BP ($2\sigma$) | Median cal yr BP |
| --- | --- | --- | --- | --- | --- | --- |
| IUACD#17C1321 | Bulk Peat | PS1 | 2-11 cm | $-159 \pm 31$ | 19 to -2 | 4 |
| IUACD#17C1322 | Bulk Peat | PS3 | 25-34 cm | $1413 \pm 34$ | 1372 to 1285 | 1320 |
| IUACD#17C1323 | Bulk Peat | PS5 | 47-56 cm | $3280 \pm 41$ | 3608 to 3401 | 3510 |
| IUACD#17C1324 | Bulk Peat | PS8 | 81-90 cm | $4202 \pm 45$ | 4852 to 4584 | 4730 |
| IUACD#17C1325 | Bulk Peat | PS10 | 104-113 cm | $6135 \pm 50$ | 7165 to 6894 | 7035 |
| IUACD#19C2461 | Bulk Peat | PS11 | 115-124 cm | $11508 \pm 49$ | 13455 to 13263 | 13355 |
| IUACD#17C1326 | Bulk Peat | PS13 | 138-147 cm | $18235 \pm 97$ | 22361 to 21846 | 22103 |

### 2.3. Macrocharcoal counts

From each of the 13 samples collected for analyses, 1 gm peat was kept in 10 ml 10% KOH solution overnight to deflocculate particles followed by NaOCl treatment to remove non-charred organic matter (Stevenson and Haberle 2005). Particles >125 μm size were collected for identification. Macrocharcoal particles are visually recognizable as opaque, angular and usually planar, black fragment and were counted using a stereomicroscope (LeicaS4E) at 10x magnification (Mooney and Tinner 2011). The extraction was performed thrice for each sample, and the mean (±1 SD) count calculated.

### 2.4. Microcharcoal, pollen count and dung fungal spore analyses

Microcharcoal and fungal spores were extracted from peat using standard methods for chemical processing of pollen (Faegri and Iversen, 1989). The procedure involves a series of acid treatments with HCl to remove carbonates, HF to remove silicate minerals and acetolysis (9:1 mixture of acetic anhydride and H2SO4) to remove polysaccharides (Erdtman, 1960). The samples were mounted on glass slides with glycerine and frequencies of microcharcoal, pollen (minimum of 800 pollen grains) and fungal spores were quantified using a microscope (Olympus CX43). Total pollen counts were done with the help of reference slides at the French Institute of Pondicherry (Tissot, Chikhi, and Nayar 1994). Black, opaque angular fragments >10 μm were identified and counted as microcharcoal particles (Clark 1988). We selected 4 common fungal taxa considered as coprophilous and semi-coprophilous, *Sporormiella spp., Sordaria spp., Delitschia spp*., and *Trichodelitschia spp*. (Perrotti and van Asperen 2019; Baker, Bhagwat, and Willis 2013) which were identified and classified at the level of genus or species based on van Geel et al. (2011); Cugny, Mazier, and Galop (2010). The ratio of coprophilous fungal spore count to total pollen count was used to estimate fungal spore abundance.

### 2.5. Lipid extraction, quantification, biomarker identification and interpretation

Total lipids were extracted from 0.2 g of the lyophilized peat samples using 9:1 v/v mixture of dichloromethane and methanol, and recovery standard consisting of 10 mg/l *n*-Hexatriacontane d_50_. The extracts were separated into two functional groups based on charge and hydrophobicity by solid phase extraction (SPE), using 6 ml glass columns packed with 500 mg of Supelco Superclean LC-NH-2 (Kim and Salem 1990). The columns were eluted with hexane (15 ml, Fraction 1: alkanes) CHCl3/isopropanol (15 ml of 2:1 v/v, Fraction 2: alkanols). The Fraction 2 was dissolved in 100 μl bis(trimethylsilyl) trifluroacetamide (BSTFA) and 100 μl of pyridine, and heated (70 °C for 120 min) to convert all the alkanols into their respective trimethylsilyl ethers.

The two different lipid fractions were quantified in an Agilent 6890N GC (Linköping University, Sweden) interfaced to an Agilent 5973 MSD mass spectrometer at 70 eV and scanned from m/z 40–600 (at 2.62 scans/s; following Ghosh, Routh, and Bhadury (2017). All eluted solvents were evaporated, Fraction 1 was re-dissolved in hexane, while Fraction 2 samples were re-dissolved in CHCl_3_: MeOH (2:1), prior to injection in splitless mode (1 μl; inlet pressure of 10 psi with a flow rate 54.3 ml/min) and separated on a HP-5 MS capillary column (5% diphenyl dimethyl polysiloxane; 30 m length, 0.25mm i.d. and 0.25 μm film thickness). All the samples were run at constant flow (1.3 ml/min) with He as carrier gas. Detection limit in the different standards ranged from 0.1 to 1 ng/g. Reproducibility of internal standards was in the range of 5-10 ppm for different compounds. All the lipid fractions were identified in samples by comparing their characteristic mass spectra, and generated ion fragments (base peak and molecular ion), retention time, and elution order published in the literature (Peters, Walters, and Moldowan 2005), NIST online library, and Archives of Mass Spectrometry.

We employed several *n*-alkane indices based on carbon chain length and ratios of different compounds to infer their origin from different plant types such as grasses, trees/shrubs, and aquatic plants. Similarly, stanols and sterols extracted from peat samples were used to infer their inputs from herbivorous mammal or human faeces (Table 2).

**Table 2.**
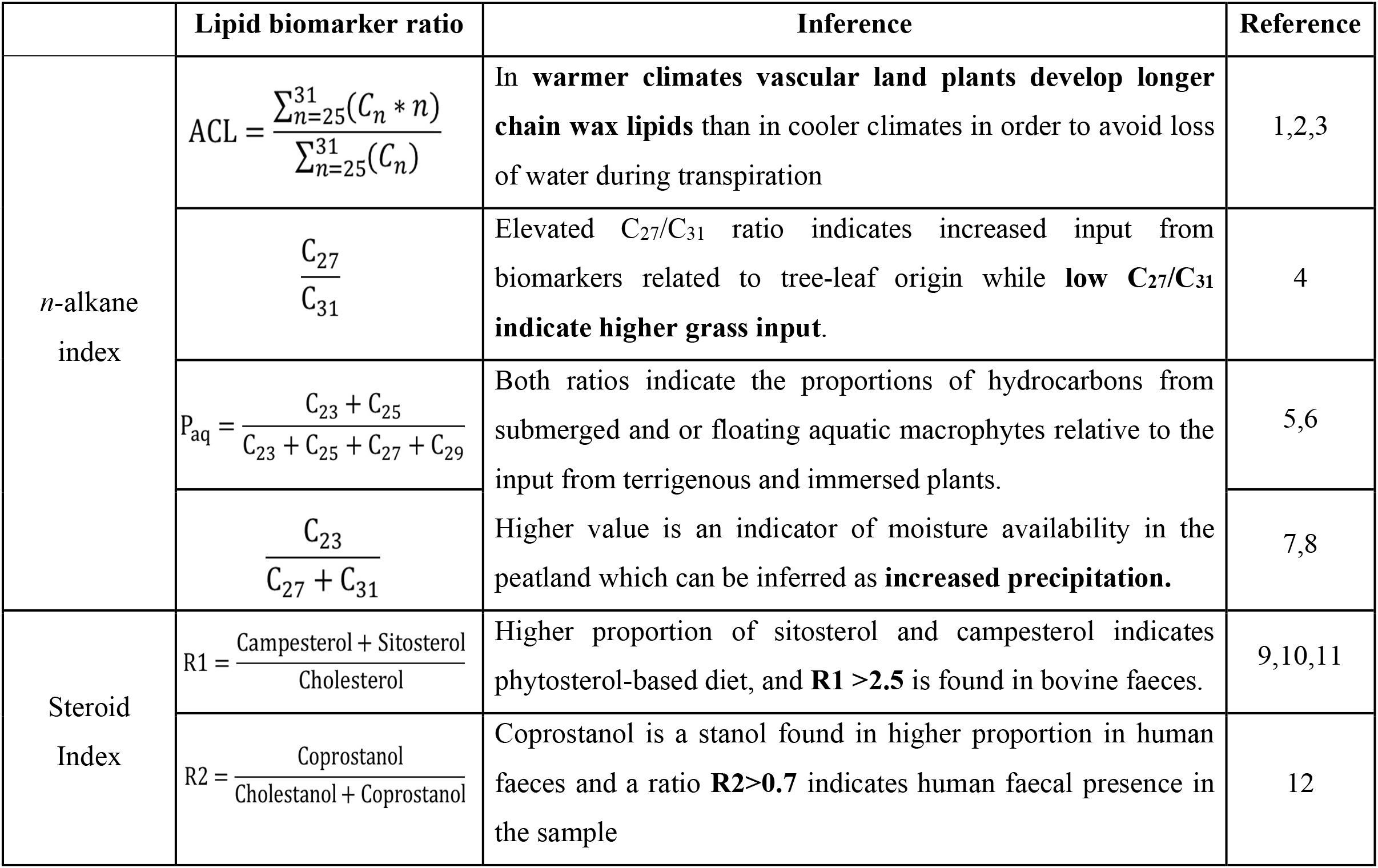
Lipid biomarker ratios used in the present study and their interpretation (References:1-Cranwell, 1973, 2-Rommerskirchen et al., 2006, 3-Gagosian and Peltzer, 1986, 4-Schwark et al 2002, 5-Ficken et al., 2000, 6-Nichols et al., 2006, 7-Andersson et al., 2011, 8-Zheng et al., 2007, 9-Jardé et al., 2007a, 10-Jardé et al., 2007b, 11-Derrien, Yang, and Hur 2017, 12-Grimalt et al., 1990)

## 3. Results

### 3.1 Chronology

AMS radiocarbon dating of peat samples from 7 depths of the Sandynallah pit were found to be in sequence without inversions; both radiocarbon and calibrated dates along with uncertainty estimates are given in Table 1. The uppermost sample in the profile near the surface (PS1) is modern in age, PS3 is dated to 1320 cal BP, and the deepest sample in the profile is dated to 22,103 cal BP. We see a faster accumulation rate in the Holocene and slower rate in Late Pleistocene sections of the profile.

### 3.2 Charcoal abundance and fire occurrence

Macrocharcoal counts from the peat samples of Sandynallah are high at ~22,000 cal BP indicative of fires; the microcharcoal counts and charcoal/pollen ratios at this time are also enhanced though to a lesser extent than the macrocharcoal counts. On the other hand, macrocharcoal shows a sharp peak ~3,500 cal BP along with corresponding peaks in microcharcoal counts and microcharcoal/pollen ratio (Fig. 2), clearly indicating intensive fire in the region at this time.

**Fig 2.**
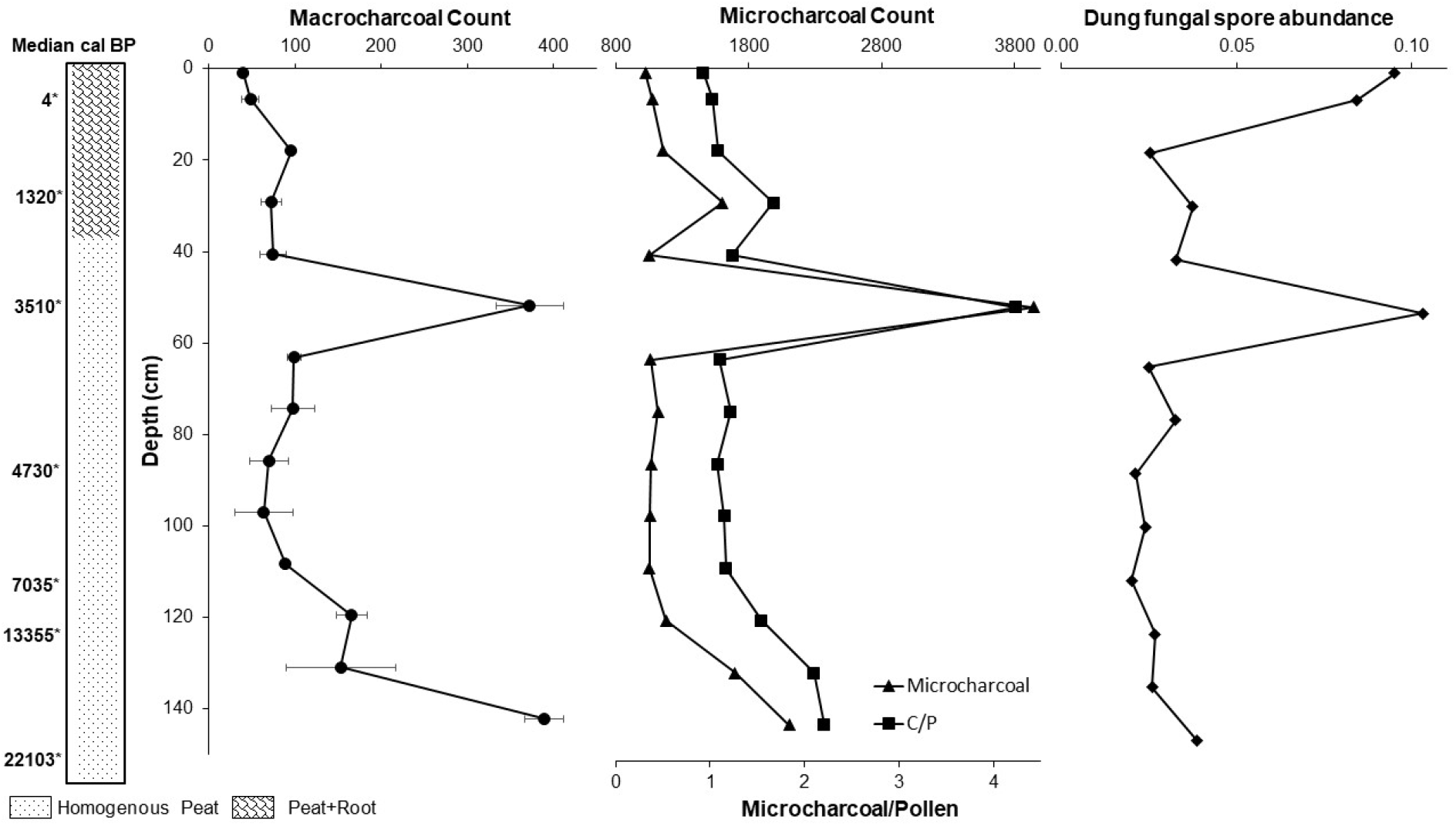
Macrocharcoal count, Microcharcoal count, Microcharcoal/Pollen ratio and the fungal spore abundance plotted against depth for the 14 peat samples at Sandynallah.

### 3.3 Dung fungal spore abundance

Coprophilous fungal spores of the taxa *Sporormiella spp., Sordaria spp., Delitschia spp*., and *Trichodelitschia spp*. were found in the peat samples at different depths. The pollen-normalized ratio of coprophilous fungal spore abundance showed two peaks, one in the modern samples and another at ~3500 cal yr BP (Fig 2). Peaks in modern samples are consistent with the increased presence of domestic livestock in the area, the peak at ~3500 cal yr BP coinciding with the peak in micro- and macrocharcoal abundances, indicating a corresponding increase in mammalian herbivore abundance around the time that intense fires were recorded.

### 3.4 Lipid biomarkers

The lipid biomarker proxies evaluated in the current study can be of three broad categories: vegetation proxies (average chain length (ACL) and C_27_/C_31_, moisture proxies (P_aq_ and C_23_+C_25_/C_23_+C_25_+ C_27_+C_29_) and steroid indices (R1 and R2) (Table 2). Total *n*-alkane content of the samples ranged from 0.12 to 22.5 μg/g of dry peat. Average chain length (ACL) in *n*-alkanes varied between 28 to 31. ACL values peaked at 22,000 cal BP and shows an increasing from 4700 cal BP onward with a peak between 3500 and 1300 cal BP. The calculated P_aq_ ranged from 0.109 to 0.954 with the 3500 cal BP sample showing the lowest value Paq value. *n*C_27_/*n*C_31_ ratio ranged from 0.13 to 5.57 in the peat samples with an average of 1.98 (Fig. 3). Changes in *n*C_23_/(*n*C_27_+*n*C_31_) over time broadly correlated with the P_aq_ value, showing two peaks at ~4700 and ~1300 cal BP. Two ratios of steroid compounds (see Table 2), one comprising sterols indicative of herbivore faecal presence and the other comprising stanols indicative of human faecal presence, from the peat samples are plotted in Figure 2. Peat sample dated at ~3500 cal yr BP showed the highest value for both ratios (R1= 20.27 and R2= 0.98) indicative of herbivore presence and human faecal presence. The sample at ~1300 cal yr BP also showed steroid ratios (R1= 5.12 and R2= 0.94) above threshold levels indicative of herbivore and human faeces. Modern samples also showed traces of herbivore faecal matter, though the steroid ratios were below the threshold of human faecal presence.

**Fig 3.**
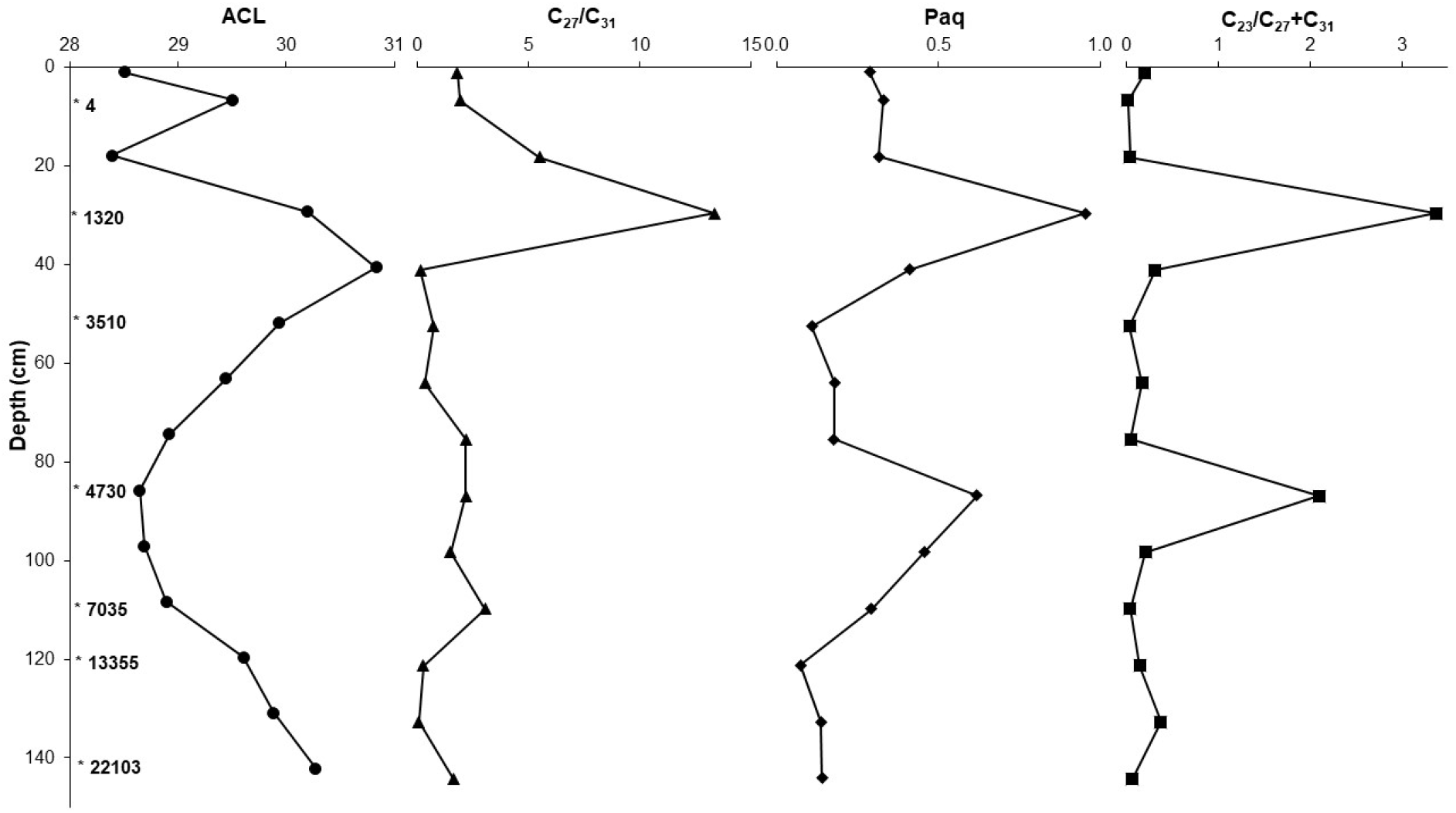
n-alkane biomarker indices, ACL and nC27/nC31 indicating vegetation shifts in the past, and Paq, nC23/(nC27+nC31) indices showing the contribution from lower chain length alkanes indicating peatland wetness (see text and Table 2 for details).

## 4. Discussion

We have shown through a combination of charcoal and pollen abundance, *n*-alkane concentration, coprophilous fungal spores and steroid lipid biomarkers in peat deposits (spanning the period ~ 22,000 cal. yr BP to the present) that intense fire activity at ~ 3500 cal. yr BP coincided with dominant grassland cover, abundance of herbivorous animals, and the presence of humans for the first time in a montane region (>2000m asl) of the Western Ghats, India. There are several questions which arise from these findings: were the fires in the extensive grasslands during this time the result of climatic desiccation or entirely due to deliberate human action?; does the abundance of herbivorous animals indicate populations of wild mammals or domesticated mammals?; who were the people inhabiting the plateau at this time?

### 4.1 Fire history from the charcoal record

The history of wildfire at a given location reflects both the prevailing climate as well as human activity and, hence, can provide valuable clues on human occupation of a site. Macrocharcoal (>125 μm) usually gets deposited close to the source of the fire making it a useful proxy for reconstructing fire history at a local scale, while microcharcoal (<125 μm) can have its origin over a much wider region, thus being a proxy for fire history over a broad spatial and temporal scale (Clark et al. 1998; Carcaillet et al. 2001; Whitlock and Larsen 2002).

Charcoal counts at Sandynallah depict two periods of enhanced fire activity, the first at ~ 22,000 cal yr BP and the second at ~3500 cal yr BP (Fig 2). The elevated levels of macrocharcoal in the sample from the deepest layer (142 cm) dated at ~22,000 cal yr BP are suggestive of burning of woody vegetation (shrubs and/or trees) (Fig 5). The sharply increased counts of microcharcoal and the high charcoal/pollen ratio clearly point to grassland fires during this period. This fire event is consistent with the expansion of C4 grasses and arid climatic conditions during the Last Glacial Maximum (LGM) recorded in paleovegetation and paleoclimate studies from the Nilgiri plateau (Sukumar et al. 1993; Caner et al. 2007). The Last Glacial Maximum dated variously across the globe at between 26-16 ka and characterized by lower temperatures and arid conditions worldwide (Clark et al. 2009; Hughes and Gibbard 2015). The Indian summer (southwest) monsoon also weakened during the LGM (Prell and Kutzbach 1987; Kumaran et al. 2013; Saraswat, Nigam, and Correge 2014) including in the northern and central Western Ghats (Sukumar et al. 1993; Rajagopalan et al., 1997; Prabhu et al. 2004; Caner et al. 2007; Kumaran et al. 2013), and this would have increased the flammability of the vegetation including forest patches in the montane Nilgiris as seen in other tropical regions (Farrera et al. 1999; Bassinot et al. 1994; Prabhu et al. 2004; Caner et al. 2007). Greater fire activity during the Last Glacial Maximum (LGM) is also observed in charcoal records from tropical latitudes of Southeast Asia (Power et al. 2008).

**Fig 5.**
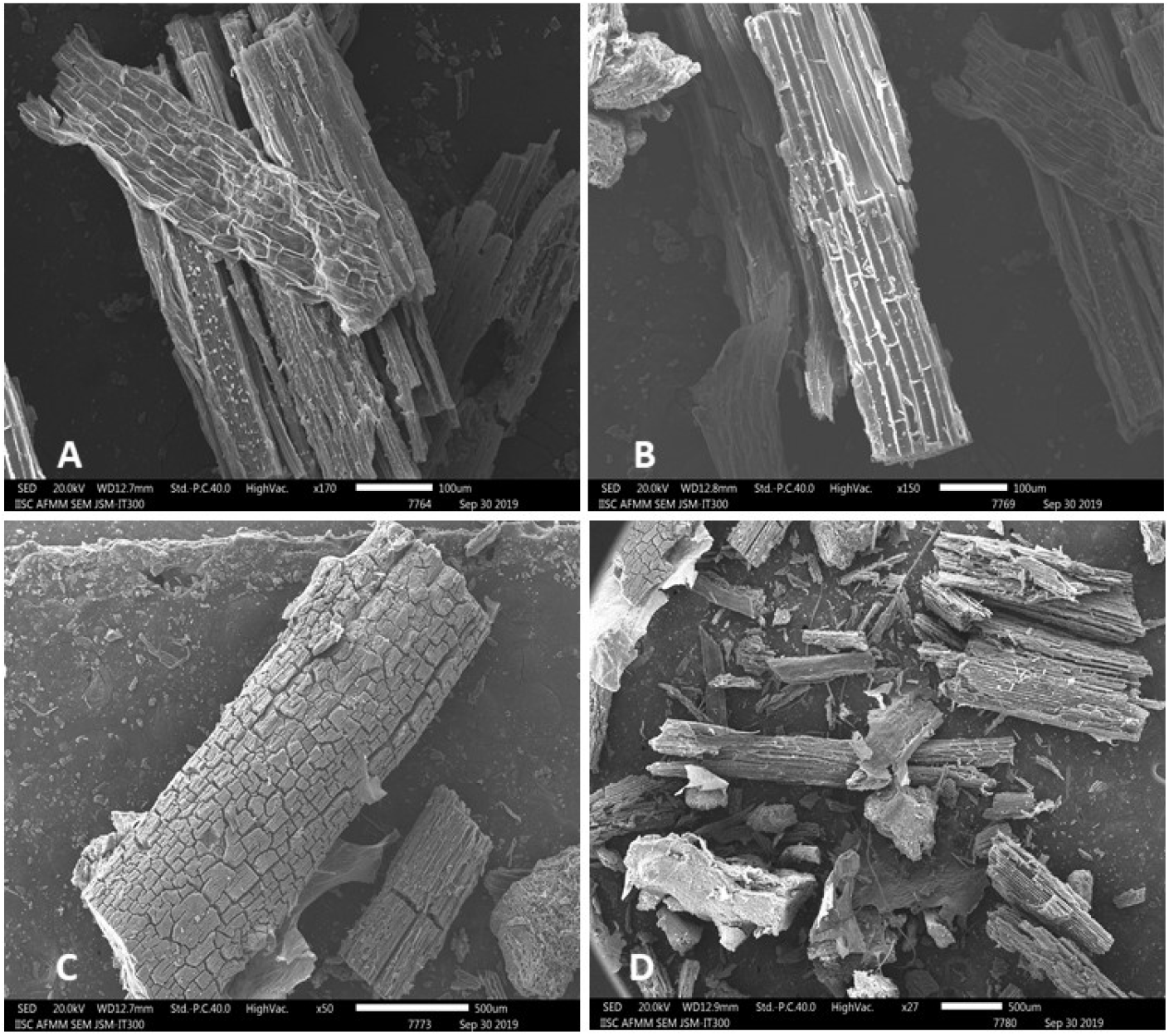
Scanning electron microscope (SEM) images of macrocharcoal particles from two fire events recorded in pit profile; A and B are particles from the 3510 cal BP peat sample, while C and D are particles isolated from the 22103 cal BP peat sample

The sharp increase in macrocharcoal counts, microcharcoal counts and C/P ratio in the peat layer from ~ 3500 cal yr BP is indicative of an intense, local fire event (Fig 2). Climate is the primary driver of wildfires globally (Flannigan et al. 2009; Krawchuk and Moritz 2011) and at local scales such as the Nilgiris (Mondal and Sukumar 2016), with successful ignition and spread of fire from human activity (as opposed to natural causes such as lightning) generally playing a secondary role (Bowman et al. 2011) It is entirely possible that an immigrant pastoralist people set fire to the natural grasslands in order to improve the pasture for their livestock, a very common practise around the world since historical times (Johansen 2004; Vuorio et al. 2014; Coughlan 2015). Paleo-ecological studies, based on stable carbon isotope ratios discriminating between C4 plants (tropical grasses) and C3 plants (trees, shrubs and other herbs), at this site have clearly pointed to an overall weakening of the monsoon resulting in expansion of grassland during ~5000-2000 cal. yr BP (Sukumar et al. 1993; Rajagopalan et al., 1997). The environmental conditions at ~3500 yr BP would have been favourable for wildfires. However, the sharp spike in both micro- and macro-charcoal abundance in the present study indicates that not only grassland but also the embedded montane forest patches burnt, thereby implying that extremely dry weather and possibly high ambient temperatures provided the conditions for an intense fire/s to spread. There would have been no necessity for a pastoralist community to set fire to forest especially when grasslands were already extensive; it is thus most likely that natural fires or anthropogenic fires in the grasslands spread to the forest patches under extremely favourable weather conditions.

Changes in pollen assemblages are used to interpret human agricultural practices, but palynological evidence from the Nilgiri region shows no indication of active cultivation in the higher elevation region except in recent times (Vasanthy 1988; Sutra, Bonnefille, and Fontugne 1997). The intense fire event ~3500 cal yr BP in the upper Nilgiri plateau featuring dominant grassland vegetation is also coincident to the abrupt transition of forest to woodland/grassland further north in the Western Ghats (North Kanara district (0-700m asl) of Karnataka state, about 500 km northwest from our montane site in the Nilgiris) at precisely the same time as evidenced in sharply enhanced counts of grass pollen and reduction of tree pollen in a marine core from the estuary of the Kalinadi River on the Arabian Sea coast (Caratini et al. 1991; Caratini et al. 1994). This remarkable match in dates of the vegetation shift in North Kanara and the large fire event in the Upper Nilgiris provides unambiguous evidence of regional aridification at ~3500 cal yr BP in peninsular India.

Increased forest clearing, landscape burning and agricultural activity after the mid-Holocene dry conditions played a crucial role in the determining the extent of vegetation cover in the mid- and lower elevations of the Western Ghats (Bhagwat et al. 2012). A study from mid-elevation (c. 900 m asl) region of Kodagu District (Karnataka State), located 170 km to the north of the Nilgiris, recorded fire activity beginning around 3500 yr BP (though peaking at a later time), lowering the tree cover state, possibly due to large-scale land use changes for settled agriculture (Bhagwat et al., 2012).

### 4.2 Lipid n-alkane biomarkers and vegetation change

The main objective of the lipid *n*-alkane biomarker analysis was to look at the vegetation history of these natural montane forest-grassland mosaics and compare this to previous paleoecological studies from the region. Organic matter accumulated in the peat consists of above-ground plant material as well as secondary products from microbial alteration and diagenesis of the primary material. We used sedimentary *n*-alkane ratios as a proxy to understand the vegetation changes in general and, more specifically, the climate associated with the fire event dating back to ~3500 cal yr BP. *n*-alkane distributions are dominated by longer chain homologues, mainly *n*-C_22-32_, indicating higher input from epicuticular leaf waxes of vascular plants and only minor contribution from algal and bacterial sources (Cranwell et al., 1987; Eglinton and Hamilton, 1967; Rieley et al., 1991; Rommerskirchen et al., 2006). Cranwell (1973) attributed elevated *n*C_31_ to grass input, whereas *n*C_27_ or *n*C_29_ predominance was related to a tree-leaf origin. Following this idea, we employed ACL and C_27_/C_31_ as vegetation indices to elucidate the grassland-forest dynamics in the past. Since this is the first attempt to interpret vegetation abundance through *n*-alkanes at Sandynallah, we are aware of the limitations of not directly accounting for relative percentage contributions of biomass from grass and woody species. However, what is striking is that in both the major fire layers, i.e. at ~22000 cal BP and ~3500 cal BP, higher ACL values and lower C_27_/C_31_ point to a higher proportion of *n*-C_31_ alkanes in the sediment, indicating a prevalence of grasslands (Cranwell, 1973; Bi et al., 2005; Rommerskirchen et al., 2006). We also considered proxies of surface wetness in the peat bog since *n*-alkane distributions of submerged and floating plants maximize at C_23_ and C_25_ while land plants maximize at C_27_, C_29_ and C_31_ (Zheng et al. 2007). The moisture indices (P_aq_ ratio and *n*C_23_/(*n*C_27_+*n*C_31_)) showed two peaks, one at ~4700 cal BP and the other at ~1300 cal BP, indicating wetter conditions; these indices however indicate very low surface wetness in the fire-impacted layers, consistent with the expectation of arid conditions during these periods (Fig 3). In past studies, the dominance of C4 plants (grasses and sedges) has been used as a proxy for dry and arid climatic conditions (Sukumar et al. 1993; Caner et al. 2007), consistent with our results of a grass-dominated, arid environment during the LGM, and the arid period at ~3500 years BP supported by fluctuating monsoon conditions during mid-Holocene (Rajagopalan et al. 1997).

### 4.3 Coprophilous fungal spores and herbivore abundance

Coprophilous fungal spores are used as a proxy for herbivore presence and abundance and to understand past pastoral activities (van Geel et al. 2011; Baker, Bhagwat, and Willis 2013; Dubois and Jacob 2016). Spores of coprophilous fungi such as *Coniochaeta, Sporormiella, Sordaria, Podospora, Delitschia* and *Trichodelitschia* are generally considered as reliable indicators of pastoralism (Cugny, Mazier, and Galop 2010; van Geel et al. 2011). The origin of the dung can be from livestock corralled at a site or from its use as manure, fuel and building materials (Linseele et al 2013). The spike in coprophilous fungal spores count at ~3500 cal yr BP in the Nilgiri plateau indicates increased presence of herbivorous animals, which could be due to introduction of livestock by immigrant people (Fig 2). At the same time, the extensive grasslands at this time, possibly expanding over several hundred years (Rajagopalan et al. 1997), could also have facilitated an increase in the population of wild herbivores. Coprophilous fungal spores alone cannot distinguish between domestic and wild fauna (Giguet-Covex et al. 2014). One mammal species which could have benefitted from grassland expansion is the Nilgiri tahr (*Nilgiritragus hylocrius*), endemic to the higher elevation grasslands of the Western Ghats, but we cannot rule out the presence of other herbivorous mammals with more widespread distribution such as the gaur (*Bos gaurus*), a bovid, and the elephant (*Elephas maximus*), found in very low numbers presently at elevations >1500 m asl. However, a 5-fold spike in coprophilous fungal spores is unlikely to reflect an upsurge in wild herbivore populations but rather a corralled herd of domestic herbivores. Currently, sheep and other livestock are maintained at Sandynallah and this is accurately picked up in the increased coprophilous fungal spore abundance in the modern samples (Fig 2).

### 4.4 Lipid steroid biomarkers and human presence

Faecal biomarkers are routinely used in archaeobotanical studies to understand anthropogenic inputs and to identify the origin of dung at a study site (Bull et al. 2001; Shillito et al. 2011; Bull et al. 2002; Bull et al. 1999). The relative ratio of two or more sterols and stanols can be used to discriminate the sources of faecal contamination (Derrien, Yang, and Hur 2017; Leeming et al. 1996). Herbivore faeces are expected to have higher proportion of sitosterol and campesterol due to their phytosterol-based diet, and the R1 >2.5 (see Table 2) is typical of faecal presence in a sample (Jardé et al.2007a; Jardé et al. 2007b; Derrien, Yang, and Hur 2017). 5α-cholestanol is the diagenetic product of cholesterol indicative of source and preservation of stanol in the environment. We used R2 ratio (Table 1) to compare the 5α-cholestanol with the coprostanol, which is a stanol found in higher proportion in human faeces to detect the presence of humans in the region (White et al. 2018; Bull et al. 2002). The 3500 cal yr. BP sample with higher R1 as well as R2 ratio indicate both human and herbivore faecal presence in the samples (Fig 4) (Derrien, Yang, and Hur 2017; Grimalt et al. 1990; Prost et al. 2017; Jardé et al. 2007a, Jardé et al. 2007b). Modern samples showed traces of herbivore faecal contamination which is expected since the study site presently maintains stocks of sheep and is also visited by cattle (Fig 4). Interestingly, both R1 and R2 ratios are above the threshold levels in the 1300-year-old sample indicating the continued presence of livestock and humans.

**Fig 4.**
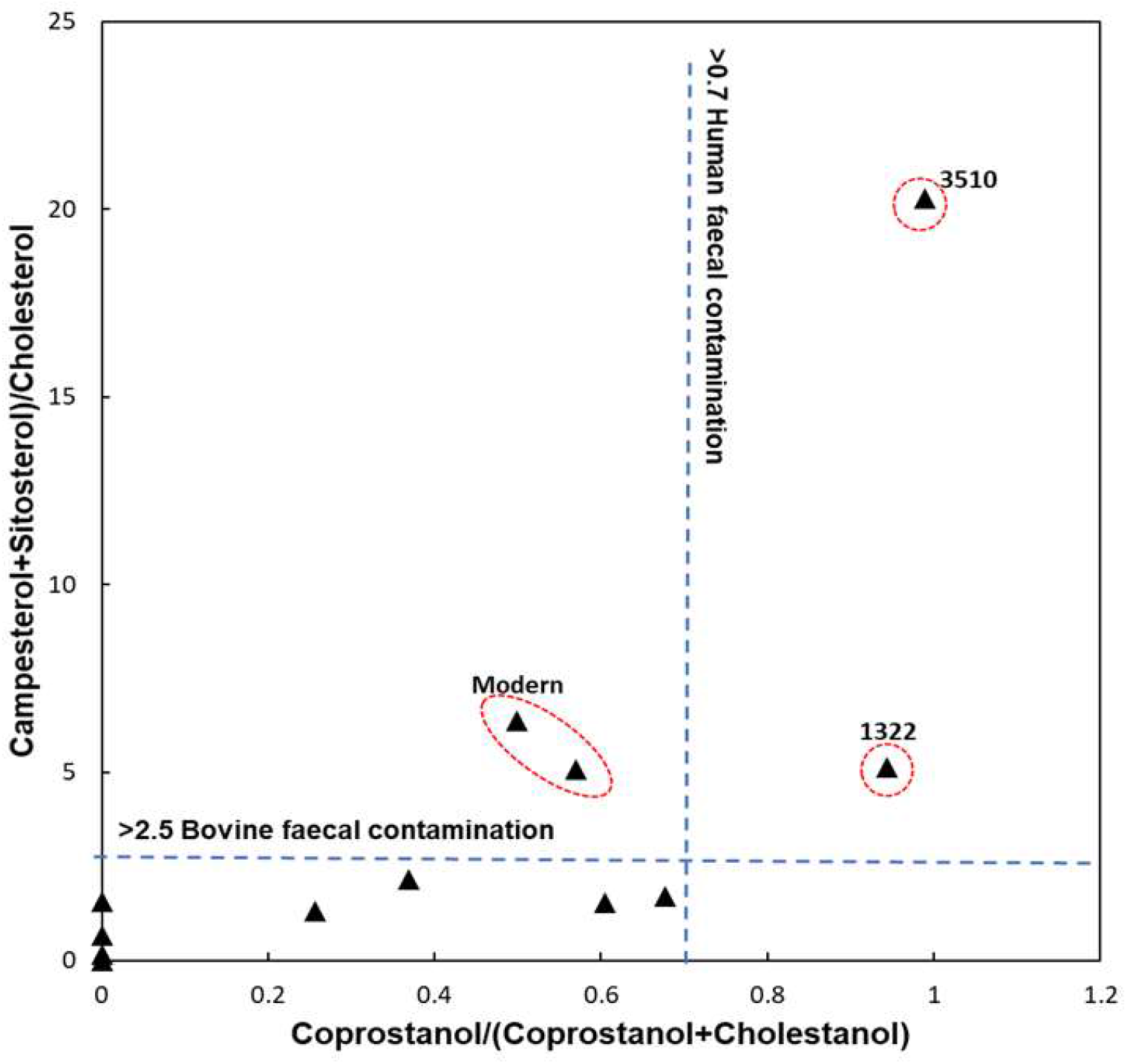
Plot of sterol biomarker ratio R1 versus R2 (see Table 2 for details), with human and bovine faecal contamination thresholds indicated by dotted lines. Biomarker ratios which are above the threshold values for bovine and human faeces are marked with a red circle along with their dates in cal BP.

### 4.5 Who were the earliest inhabitants of the upper Nilgiri plateau?

There are five traditional peoples or “tribes” in the Nilgiri hills – the Kurumbas, the Irulas, the Badagas, the Kotas and the Todas (Noble 1967).(Noble 1967) Of these, the Kurumbas and the Irulas inhabit the lower elevations (1600-600m asl) of the Nilgiris, practise subsistence agriculture or are hunter-gatherers and, thus, are unlikely to be the earliest inhabitants of the upper Nilgiri plateau. The Badagas are mainly cultivators inhabiting the upper plateau but are known to have migrated to the Nilgiris from the northern Karnataka region only during the late 16th century CE following the defeat of the Vijayanagar Empire (Francis 1908; Hockings 1980). The Kotas in the upper plateau are essentially artisans, musicians, and cultivators, though, like the Badagas, they may also tend some buffalo presently.

The most obvious candidates for the earliest inhabitants would be the Todas, a near-obligate pastoralist people believed to have inhabited the Nilgiri plateau around the 1st century CE and remained relatively isolated to the outside world until they were “discovered” during the early 19th century by the erstwhile British administration (Rivers 1906; Emeneau 1997; Walker 1997). Local legend also generally concedes that the Todas are the original inhabitants of the Nilgiri plateau. Based mainly on linguistic affinities of the Toda language with the common Dravidian languages of southern India, it has been suggested or even assumed that the Toda origin in the Nilgiris is not more than 2000 years old (Rivers 1906; Noble 1976; Emeneau 1997; Zagarell 1997). Our study provides compelling evidence for the presence of people and livestock at Sandynallah in the higher elevations of the Nilgiris at 3500 cal BP coinciding with or subsequent to a changed climate and environment in peninsular India. This pastoralist people also probably managed the landscape through setting grassland fires whose spread was aided by favourable weather conditions. The most parsimonious interpretation would be that these people were indeed the Todas who had immigrated along with their buffaloes to this location by 3500 cal BP, or more than 1500 years prior to what has been believed so far. Genetic studies suggest that the Toda buffalo is most closely related to the buffaloes further north along the Western Ghats in the South Kanara district of Karnataka state (Kathiravan et al. 2011). It is of course entirely possible that the people and livestock recorded at 3500 cal BP represent an unknown tribe or even a failed immigration. In any case, the evidence opens up the interdisciplinary fields of paleoecology, archaeology, and human ecology during mid-to late Holocene for further investigation not only in the montane Nilgiris but the broader region of peninsular India.

## Acknowledgments

RS was a JC Bose National Fellow during the tenure of this study. DG thanks DST INSPIRE Faculty Funds (Grant No. DST/INSPIRE/04/2015/002362) for lipid biomarker analysis. SPK thanks Indian Institute of Science, Bangalore, and Inspire DST for the MS fellowship. We thank Dr. Anupama Krishnamurthy, Mr. Prasad S, and Mr. Orukaimani of French Institute, Puducherry who provided support and guidance in pollen slide preparation and valuable insights for the project. PRB thanks Prof. Iyue, Tamil Nadu Veterinary and Animal Sciences University (TANUVAS) and the staff at Sheep Breeding Research Station (SBRS), Sandynallah for permission and assistance with sample collection. Authors are thankful to IUAC for extending graphitization laboratory and AMS measurement facility for ^14^C established under Ministry of Earth Science (MoES), Govt of India funded project with ref. no. MoES/16/07/11(i)-RDEAS.

